# Automated Incorporation of Pairwise Dependency in Transcription Factor Binding Site Prediction Using Dinucleotide Weight Tensors

**DOI:** 10.1101/078212

**Authors:** Saeed Omidi, Mihaela Zavolan, Mikhail Pachkov, Jeremie Breda, Severin Berger, Erik van Nimwegen

## Abstract

Gene regulatory networks are ultimately encoded by the sequence-specific binding of (TFs) to short DNA segments. Although it is customary to represent the binding specificity of a TF by a position-specific weight matrix (PSWM), which assumes each position within a site contributes independently to the overall binding affinity, evidence has been accumulating that there can be significant dependencies between positions. Unfortunately, methodological challenges have so far hindered the development of a practical and generally-accepted extension of the PSWM model. On the one hand, simple models that only consider dependencies between nearest-neighbor positions are easy to use in practice, but fail to account for the distal dependencies that are observed in the data. On the other hand, models that allow for arbitrary dependencies are prone to overfitting, requiring regularization schemes that are difficult to use in practice for non-experts.

Here we present a new regulatory motif model, called dinucleotide weight tensor (DWT), that incorporates arbitrary pairwise dependencies between positions in binding sites, rigorously from first principles, and free from tunable parameters. We demonstrate the power of the method on a large set of ChIP-seq data-sets, showing that DWTs outperform both PSWMs and motif models that only incorporate nearest-neighbor dependencies. We also demonstrate that DWTs outperform two previously proposed methods. Finally, we show that DWTs inferred from ChIP-seq data also outperform PSWMs on HT-SELEX data for the same TF, suggesting that DWTs capture inherent biophysical properties of the interactions between the DNA binding domains of TFs and their binding sites.

We make a suite of DWT tools available at dwt.unibas.ch, that allow users to automatically perform ‘motif finding’, i.e. the inference of DWT motifs from a set of sequences, binding site prediction with DWTs, and visualization of DWT ‘dilogo’ motifs.

**Author Summary:** Gene regulatory networks are ultimately encoded in constellations of short binding sites in the DNA and RNA that are recognized by regulatory factors such as transcription factors (TFs). For several decades, computational analysis of regulatory networks has relied on a model of TF sequence-specificity, the position-specific weight-matrix (PSWM), that assumes different positions in a binding site contribute independently to the total binding energy of the TF. However, in recent years evidence has been accumulating that, at least for some TFs, this assumption does not hold. Here we present a new model for the sequence-specificity of TFs, the dinucleotide weight tensor (DWT), that takes arbitrary dependencies between positions in binding sites into account and show that it consistently outperforms PSWMs on high-throughput datasets on TF binding. Moreover, in contrast to previous approaches, DWTs are directly derived from first principles within a Bayesian framework, and contain no tunable parameters. This allows them to be easily applied in practice and we make a suite of tools available for computational analysis with DWTs.

## Introduction

Gene regulatory networks are a crucial component of essentially all forms of life, allowing organisms to respond and adapt to their environment, and allowing multi-cellular organisms to express a single genotype into many different cellular phenotypes. Transcription factors (TFs) are central players in gene regulatory networks that bind to DNA in a sequence-specific manner. Although the molecular mechanisms through which TFs regulate expression of their target genes involve a complex interplay of interactions between TFs, co-factors, chromatin modifiers, and signaling molecules, gene regulatory networks are ultimately genetically encoded by constellations of transcription factor binding sites (TFBSs) to which the TFs bind in a sequence-specific manner.

Consequently, a key question in the analysis of gene regulatory networks is to find a proper mathematical representation of the sequence-specificities of TFs. That is, for each TF, we want to determine an energy function *E*(*s*) that calculates, for any given DNA segment *s*, the binding free energy of the TF binding to *s*. The segment *s* is generally of fixed length for a given TF, which typically ranges from 6 to 30 base pairs. Although there have been some attempts to use direct structural and biophysical modeling of the sequence-specificity of TFs, e.g. [1–3], such efforts have generally achieved only limited accuracy. Instead, by far the most common approach to representing the sequence-specificity of TFs is through a statistical mechanical analysis, which essentially assumes that the probability that a binding site for a particular TF has sequence *s* is given by a maximum entropy distribution with respect to its binding energy *E*(*s*), i.e. *P*(*s*) ∝ *e*^*λE*(*s*)^ [4, 5]. Using this assumption, the binding energies *E*(*s*) of sequence segments *s* can in principle be inferred from data on the relative frequencies *P*(*s*) with which different sequences *s* are bound by a given TF. However, the number of possible sequence segments *s* is 4^*l*^, which is already over a million for relatively short TFBSs of length *l* = 10 base pairs, i.e. much larger than the total number of genome-wide binding sites for a single TF. Thus, a crucial additional assumption, that has been made for several decades [6], is to assume that each base pair in the binding site contributes *independently* to the overall binding energy, i.e. 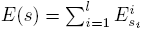,where *s*_*i*_ is the base occurring at position *i* in sequence segment *s* and 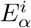 is the energy contribution of base *α* at position *i*. Under this independence assumption, the sequence-specificity of a TF can be parametrized by 3 *× l* parameters

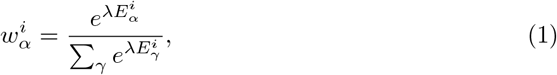

where 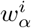 is the fraction of binding sites that have letter *α* at position *i*. This is the well-known position specific weight matrix (PSWM) representation which has been used in the vast majority of works on modeling TF binding and TFBS prediction. The main advantage of this approach is the relatively small number of parameters, allowing reasonable estimation of the weight matrix entries 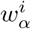 from as few as a dozen of example binding sites.

With the drastic reduction in costs of DNA sequencing over the last decade and the development of a number of experimental techniques for identifying TFBSs in high-throughput, such as ChIP-seq [7], protein binding arrays [8], and HT-SELEX [9], hundreds if not thousands of example TFBSs for a single TF can now be routinely obtained. Such large collections of TFBSs have enabled researchers to investigate to what extent the assumption of independence, i.e. that each position in the binding site contributes to the binding energy independent of the other positions, holds in practice. The results of these investigations indicate that, although the assumption of independence is often reasonably accurate, there are also many cases which clearly deviate from independence.

Studies going back over a decade, such as [10] and [11], had already provided evidence that PSWMs can be unsatisfactory in describing DNA binding specificities of particular TFs, and that the assumption of independence often breaks down. More recently, a large-scale study by Bulyk and colleagues assayed 104 distinct mouse TFs using protein binding microarray (PBM) technology and found that, for a large fraction of the TFs investigated, the binding energy landscapes were significantly more complex than assumed by PSWM models [12]. Notably, a number of assayed TFs exhibited strong support for pairwise dependencies (PDs) within their binding sites. As another example, Nutiu *et al.* [13] studied the binding specificity of the yeast TF Gcn4p in detail and showed that it exhibits several strong PDs. Moreover, a model that incorporates these PDs was shown to outperform PSWM models in explaining the observed TFBSs. In summary, all these results suggest that accurate representation of TF sequence-specificities requires that dependencies between positions are taken into account, although it remains unclear how important such dependencies are for the accuracy of TFBS prediction.

## Incorporating pairwise dependencies

Several works have modeled TF binding specificity by including dependence between binding positions. A major challenge is that, when an arbitrary number of dependencies between arbitrary pairs of positions is allowed, the number of possible models and parameters grows rapidly, so that it becomes difficult to reliably identify the best models, and to avoid overfitting. Previous works have taken different approaches for addressing this challenge.

In some approaches, model complexity is directly controlled by only allowing dependencies between adjacent positions, e.g. [14, 15]. However, previous analyses indicated that substantial dependencies can occur between more distal pairs of positions, and our analyses below also indicate that significant dependencies between non-neighboring positions are common. Below we explicitly compare our general DWT model with a restricted model that only incorporates dependencies between adjacent positions.

In other approaches, PDs between arbitrary pairs of positions are in principle allowed, but instead of incorporating all possible pairwise dependencies, different *ad hoc* approaches are employed to restrict the number of PDs that are taken into account. For example, a Bayesian network model by Barash *et al.* [16] starts by calculating likelihoods for all possible PDs, finds the spanning tree of PDs that has maximum likelihood (ML), and then models the TF binding specificity using only the PDs in this ML spanning tree. That is, of the *l*(*l* − 1)/2 possible PDs, only (*l* − 1) end up being used for modeling the TF binding specificity. As another example, the variable-order Bayesian network model of Grosse and Grau [17] starts from a full higher order Markov model (represented as a tree of possible sequence contexts) and then reduces the number of parameters by systematically collapsing different sequence contexts that do not show significantly different statistics in the data, i.e. pruning the tree.

Alternatively, some approaches start from a model without dependencies, and use a greedy algorithm that iteratively adds PDs which maximally improve the model. For example, Sharon *et al.* [18] express the TF’s binding specificity as a weighted sum of features, where features are propositions that can either be true or false, e.g. a specific pair of nucleotides appears at a particular pair of positions. Features are iteratively added to the model until no additional feature can be found that further improves the model. However, this iterative procedure often leads to overfitting and Sharon *et al.* used a combination of regularization procedures to control model complexity.

A similar iterative approach is used in the work of Santolini *et al.* [19] where the TF binding specificity is modeled by an inhomogeneous Potts model, which incorporates information from both single and pairs of positions. Individual pairs of positions are iteratively added to the model so as to maximize its likelihood. Here too the authors find that this procedure can easily lead to overfitting and they use the Bayesian information criterion as a regularization scheme to penalize model complexity. Below we will compare the performance of our approach with both the approaches of Sharon *et al.* [18] and Santolini *et al.* [19].

In spite of these efforts, no model that incorporates PDs has found widespread application in the community so far. Models that only use adjacent positions are attractive for their simplicity, but fail to capture the distal PDs that are clearly evident in the data. In contrast, models that consider arbitrary PDs make use of *ad hoc* approaches to restrict the number of PDs considered, and employ complex regularization schemes that require expert supervision, which make them harder to use in practice. The current challenge is thus to develop a model that, on the one hand, rigorously incorporates all possible PDs, and that is easy to use in practice, i.e. not requiring expert tuning of parameters or control of model complexity, on the other hand.

Here we present a new Bayesian network model, called dinucleotide weight tensor (DWT), which takes into account all possible PDs within a rigorous probabilistic framework that has no tunable parameters and automatically avoids over-fitting. In particular, in the DWT model all unknown parameters including the topology of the network of direct interactions and the joint probabilities for all dependent pairs of nucleotides within the network are analytically marginalized over, so that binding energies *E*(*s*) that take all PDs into account can be calculated from first principles, and without the need for the user to set any tunable parameters. This makes the DWT model highly robust and easily applicable in practice, i.e. even when there are no significant PDs. Indeed, in addition to presenting the algorithm below, we have also developed a suite of software tools that can be used to perform motif finding with DWTs, visualization of DWT motifs, and TFBS prediction with DWTs, which we make publically available with this publication.

We demonstrate the power of the DWT approach using a large collection of 121 ChIP-seq data-sets representing 92 different human TFs. We show that DWTs outperform PSWMs for a substantial fraction of the TFs, and never perform substantially worse, demonstrating that DWTs automatically avoid over-fitting, even though there are no explicit regularization schemes. Second, we show that DWTs outperform a restricted model that only incorporates dependencies between adjacent positions for the large majority of datasets, demonstrating that distal positions contribute to the accuracy of TFBS prediction. We also show that DWTs substantially outperform two previous approaches [18, 19]. Finally, using HT-SELEX data for a set of TFs for which ChIP-seq data are also available, we show that the DWTs inferred from ChIP-seq data also generally outcompete PSWMs on HT-SELEX data. Since the HT-SELEX experiments are performed *in vitro* using only the DNA binding domains of the TFs, these results suggest that the DWT likely captures aspects of the biophysical interaction between the DNA binding domains of the TFs and their cognate binding sites.

## Materials and Methods

### The Dinucleotide Weight Tensor model

We here present the dinucleotide weight tensor (DWT) model for describing TF sequence-specificities using arbitrary pairwise dependencies. The DWT model is based on a Bayesian network model that we have applied previously to model interactions between proteins [20] and to predict contacting residues within three-dimensional protein structures [21]. The model describes the probability distribution *P*(*s*) of binding site sequence segments *s* as a mixture of all possible factorizations of the joint distribution over *s* into pairwise conditional probabilities between pairs of positions in *s*.

Let *S* denote an ungapped alignment of sequences of a given length *l*, that are hypothesized to correspond to a collection of binding sites for a common TF. A central quantity in probabilistic motif finding is the probability *P*(*S*) that this collection of sequences derives from a common PSWM *w*. Under the assumption of independence that the PSWM model makes, the probability *P*(*S*) is given by a product of the probabilities *P*(*S*_*i*_) for the individual alignment columns *S*_*i*_, i.e. 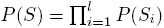.Formally, the probability *P*(*S*_*i*_) is given by an integral over all possible PSWM columns 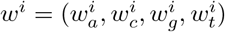, i.e. *P*(*S*_*i*_) = ∫ *dw*^*i*^*P*(*S*_*i*_|*w*^*i*^)*P*(*w*^*i*^), where *P*(*w*^*i*^) is a prior probability density on the PSWM column and the integral is over the simplex 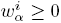, 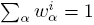. Using a Dirichlet prior of the form 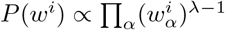, the integral can be performed analytically and yields

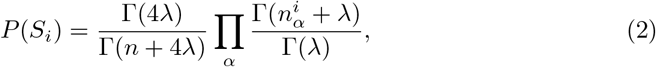

where 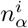 is the number of sequences in *S* that have letter *α* at position *i*, *n* is the total number of sequences in *S*, and Γ(*x*) is the gamma-function, see e.g. [5].

Here we generalize the PSWM model by assuming that arbitrary pairwise dependencies can occur between pairs of positions. In complete analogy with the calculations for the PSWM above, we can introduce a dinucleotide weight tensor *w* for the pairs of positions (*i, j*), with components 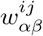 denoting the probability that the combination of letters (*α, β*) occurs at the positions (*i, j*). Using a Dirichlet prior 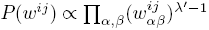 and integrating over all possible *w*^*ij*^ we then obtain the probability *P*(*S*_*i*_*, S*_*j*_) for a pair of columns (*i, j*) in complete analogy with the PSWM case

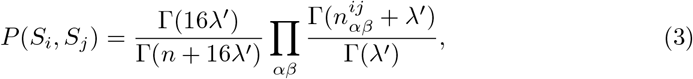

where 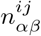 is the number of times the combination of letters (*α, β*) appears at the pair of positions (*i, j*). As we explained previously [20] consistency of the mono- and di-nucleotide priors requires that *λ* = 4*λ′*. As for the PSWM case [22], the results are generally insensitive to the precise setting of 0 *< λ ≤* 1 and we use the Jeffrey’s prior *λ* = 1/2 throughout.

The evidence for dependency in the frequencies of letters at positions (*i, j*) can be quantified by the likelihood ratio *R*_*ij*_:

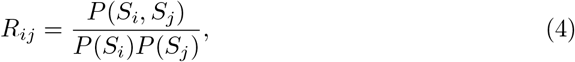

and as we will see below, the matrix *R* of these dependencies *R*_*ij*_ will play a crucial role in the calculations. As a side remark on the interpretation of the dependencies *R*_*ij*_, in the limit of a large number of sequences *n*, the Gamma-functions are well approximated by the Stirling approximation Γ(*x* + 1) *≈ x^x^* exp(−*x*) and using this it is easy to show that *R*_*ij*_ ≈ *e*^*nIij*^, where *I*_*ij*_ is the mutual information of the letter frequencies in columns *i* and *j*.

In contrast to the PSWM model, we do not assume that the probability *P*(*S*) simply factorizes into independent probabilities *P*(*S*_*i*_) for each column *i*. Instead, we will approximate the joint probability *P*(*S*) as a mixture of all possible factorizations into pairwise conditional probabilities of the form *P*(*S*_*i*_*|S*_*j*_)*P*(*S*_*j*_*|S*_*k*_)*P*(*S*_*k*_*|S*_*m*_) *…*. For any such factorization, there is a single ‘root’ position that is not dependent on any other position, and each other position *i* is dependent on one ‘parent’ position *π*(*i*). If we consider each position *i* a node of a graph, and draw an edge between each node and its parent node *π*(*i*), then each possible factorization *π* corresponds to a spanning tree of the set of *l* nodes. Noting that the conditional probability *P*(*S*_*i*_*|S*_*j*_) of column *i* given column *j* can be written as *P*(*S*_*i*_*|S*_*j*_) = *R*_*ij*_*P*(*S*_*i*_), we obtain for the probability *P*(*S|π*) of the alignment given a particular factorization *π*:

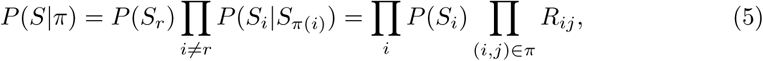

where *r* is the root node and the product on the right-hand side is over all edges in the spanning tree *π*. Note that the first product on the right-hand side corresponds precisely to the probability *P*(*S*) under the PSWM model of equation (2). The product over the dependencies *R*_*ij*_ along the edges (*i, j*) of the spanning tree *π* thus precisely quantifies the effects of the pairwise dependencies.

Instead of assuming one particular factorization *π*, we consider all possible factorizations and explicitly marginalize over the unknown factorization. That is, we aim to calculate

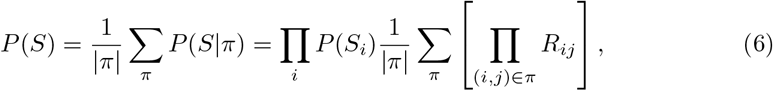

where *|π|* = *l*^*l*−2^ is the number of spanning trees of a complete graph with *l* nodes. To calculate *P*(*S*) we thus need to sum the product of the *R*_*ij*_ over all edges in the spanning tree *π* over all possible spanning trees, which may seem intractable given the large number of possible spanning trees. However, using a generalization of Kirchhoff’s matrix-tree theorem, this sum can be calculated efficiently as the determinant of an *l* − 1 by *l* − 1 matrix [20, 21, 23].

Specifically, the Laplacian *L*(*R*) of matrix *R* is obtained by replacing, for each row *i*, the diagonal element *R*_*ii*_ = 0 with minus the sum of the entries on the row, i.e. *L*(*R*)_*ii*_ = − ∑_*j ≠ i*_ *R*_*ij*_, and *L*(*R*)_*ij*_ = *R*_*ij*_ when *i* ≠ *j*. If we define *D*(*R*) to be any minor of the Laplacian *L*(*R*) of matrix *R*, we finally obtain

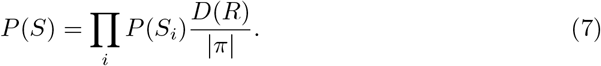

The determinant *D*(*R*) can be calculated efficiently, i.e. in *O*(*l*^3^) steps. One complication in practice is that, when there are many sequences in *S* and strong dependencies between some positions, the elements of *R* may vary over many orders of magnitude, causing the numerical calculation of the determinant to become unstable. In the supporting information we describe how we control numerical stability using a rescaling procedure.

### Binding site prediction with DWTs

We first briefly review binding site prediction using PSWMs. Assume a set of known TFBSs *S* for a particular TF is given. To predict new TFBSs for this TF one calculates the probabilities *P*(*s|S*) that, sampling another sequence from the same PSWM that the set *S* derives from, one would obtain sequence segment *s*. This probability is given by the ratio of the probability *P*(*s, S*) that all sequences derive from a common PSWM and the probability *P*(*S*) that the sequences in *S* derive from a common PSWM. Using equation (2) we have

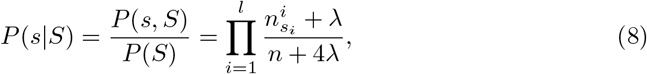

where 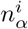 is the number of times letter *α* occurs at position *i* in the set *S*, and *s*_*i*_ is the letter at position *i* in sequence *s*. As the probabilities *P*(*s|S*) only depend on the base counts 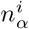, a PSWM is specified by specifying these counts (and the parameter *λ* of the prior), and the probability to sample any other sequence segment *s* from this PSWM is then given by (8).

These calculations generalize in a straight-forward manner to our DWT model. The probability to sample sequence segment *s* from the same DWT model as the set *S* is given by

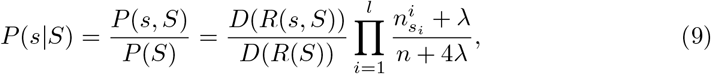

where *R*(*s, S*) is the dependency matrix *R* obtained from the full set of sequences (*s, S*) and *R*(*S*) is the dependency matrix obtained from the set of sequences *S*. Equation (9) nicely illustrates that the probability *P*(*s|S*) is given by a product of two factors: The first is identical to the PSWM model’s probability, and the second, which incorporates the effects of the dependencies, is given by a ratio of two determinants. As we will see below, for TFs where there are no significant dependencies, the latter ratio automatically becomes 1 and the DWT model automatically reduces to the PSWM model.

Whereas the probabilities *P*(*s|S*) for the PSWM model depend only on the counts 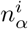, for the DWT model the probabilities *P*(*s|S*) depend on the pair counts 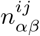. Thus, instead of specifying a set of binding sites *S*, we specify a DWT model *M* by the set of 16*l*(*l* − 1)/2 counts 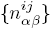 and calculate the probabilities *P*(*s|M*) using equation (9).

Finally, as explained in the supporting information, we adapted the rescaling procedure explained above to ensure numerical stability of the ratio of determinants in (9) while at the same time guaranteeing that *P*(*s|S*) remains exactly normalized, i.e. ∑_*s*_ *P*(*s|S*) = 1 when summing over all possible length-*l* sequences *s*.

### Motif finding with DWTs

To infer a motif *M* from a given set of input sequences, we need to define the likelihood function, i.e. the probability of observing our set of input sequences given the motif model *M*. Whether our input sequences derive from ChIP-seq, HT-SELEX, or a similar experimental procedure, what distinguishes the input sequences from other sequences is that they were bound by the TF in question. Thus, the likelihood should reflect the probability that the observed input sequences were bound to the TF, whereas typical ‘random’ sequences were not. We formalize this idea by imagining that we have a very large set of sequences, with nucleotide composition according to some background model, and that we are sampling sequences from this set in proportion to the probability that they are bound by the TF. The likelihood of our data set of input sequences *S* is then the probability to sample these input sequences from the large pool.

Specifically, we will assume TF binding is well approximated by a thermodynamic equilibrium model and define, for any length-*l* sequence segment *s* its effective ‘binding energy’ (in units of *kT*) as

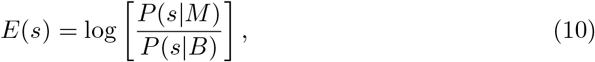

where *P*(*s|M*) is calculated as described in the previous section and *P*(*s|B*) is the probability of the sequence segment *s* under a background model. In this study we use a simple single nucleotide background model, i.e. 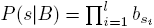 with *b*_*α*_ the overall frequency of letter *α* in the input data. Under a simple thermodynamic model, the probability *P*_*b*_(*s|M, c, E*_0_) that an isolated sequence segment *s* is bound by the TF is given by

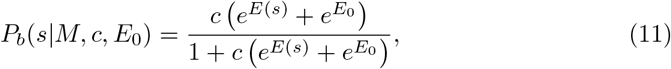

where *c* is the concentration of the TF and *E*_0_ is the energy with which the TF can be bound to *s* in a non-specific (i.e. sequence independent) manner. Note that the constant 1 in the denominator corresponds to the statistical weight of the unbound state. In this work we will assume that the concentration *c* of the TF is sufficiently small that binding is not saturated at any of the sequence segments. In this limit, the denominator can be ignored and the probability of binding is well approximated by *P*_*b*_(*s|M, c, E*_0_) *≈ c (e*^*E*_(*s*)_^ + *e*^*E*_0_^). Moreover, for a longer sequence *S*, the binding probability *P*_*b*_(*S|M, c, E*_0_) is just the sum of the binding probabilities at each of the segments of *S*:

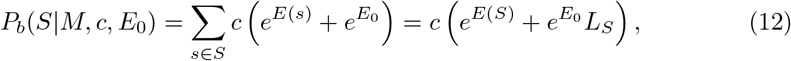

where we have defined the total binding energy *E*(*S*) of a longer sequence *S* as

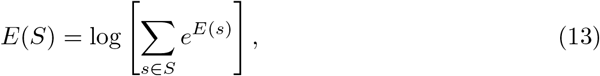

and *L*_*S*_ is the number of sequence segments in *S*, which includes segments on both the positive and negative strand of the sequence *S*.

For a large set of sequence segments sampled from the background distribution *P*(*s|B*), the average binding probability is given by

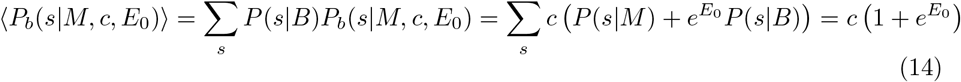

Thus, the total amount of binding to a large set of *B* background segments is *Bc* (1 + *e*^*E*_0_^) and, consequently, the probability to sample the sequence *S* from this large pool of sequences is given by

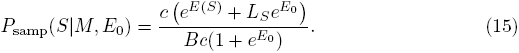

Note that the concentration *c* cancels from this expression, i.e. in the limit that binding is not saturated the relative amounts of binding to different sequences becomes independent of the precise concentration of the TF.

Finally, our desired log-likelihood *L*(*M, E*_0_) is the log-probability to sample all the sequences *S* from our input dataset *D* from the large pool of background sequences:

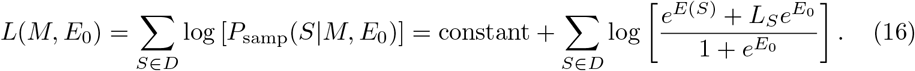

The aim of the motif finding is to maximize this log-likelihood. To do this our algorithm starts from an initial PSWM model *w* and uses an expectation maximization (EM) algorithm analogous to those used for inferring PSWMs [24] to iteratively improve *L*(*M, E*_0_).

We initialize the DWT from a PSWM that can either be specified by the user, e.g. when a known PSWM motif is already available for the TF in question, or it can be obtained by running a standard PSWM motif finder on the input sequences *S*. The sequences in the set *S* are generally longer than the length *l* of the motif but typically not longer than a few hundred base pairs, e.g. they could consist of the binding peaks obtained in a ChIP-seq experiment.

We then iterate the following steps. First we calculate the binding energies *E*(*s*) for each of the length-*l* segments in the input sequences *S*, and the total binding energies *E*(*S*) of each input sequence. Second, we optimize the non-specific binding energy *E*_0_ by finding the root of

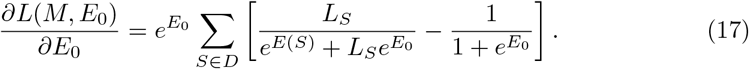

Third, we predict binding sites in the sequences *S* to calculate the pair counts 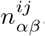 i.e. the expected number of binding sites that have the pair of letters (*α, β*) at positions (*i, j*). In particular, the probability that the TF is bound in a sequence-specific manner to sequence *S* is

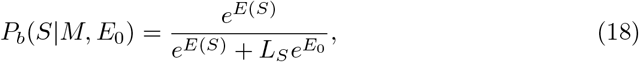

and the probability that it is bound at the specific segment *s* is

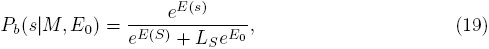

The updated pair counts 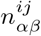 are then simply given by summing the binding probabilities *P*_*b*_(*s|M, E*_0_) over all sites in which letters (*α, β*) occur at the positions (*i, j*). These updated pair counts then define the DWT motif *M* for the next iteration, and this procedure is iterated until convergence.

### Dilogos graphically represent DWT models

To visualize DWT models, we propose a graphical representation which generalizes the well-known sequence logo and which we call a ‘dilogo’. As an example, Fig 1 shows the dilogo for the TF NRF1, which we constructed from ENCODE ChIP-seq data (see below).

The dilogo first of all shows the classical sequence logo representation of the marginal probabilities 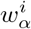 at the top. For example, in this example positions 3 − 5 are most likely to show the pattern CGC. Secondly, at the bottom the dilogo shows information about pairwise dependencies evident in the DWT. As explained in the supplementary materials and in previous work on protein contacts [25], we can calculate for each pair of positions (*i, j*) the posterior probability *P*(*i, j*) that the factorization of *P*(*S*) contains a direct dependence between positions *i* and *j*. The probabilities *P*(*i, j*) are shown in a square lattice, with the intensity of the color corresponding to the posterior probability. For example, for NRF1 there are high posterior probabilities of interaction between positions (2, 3), (2, 4), (2, 5), (6, 7), (6, 10), and (7, 8).

**Fig 1.**
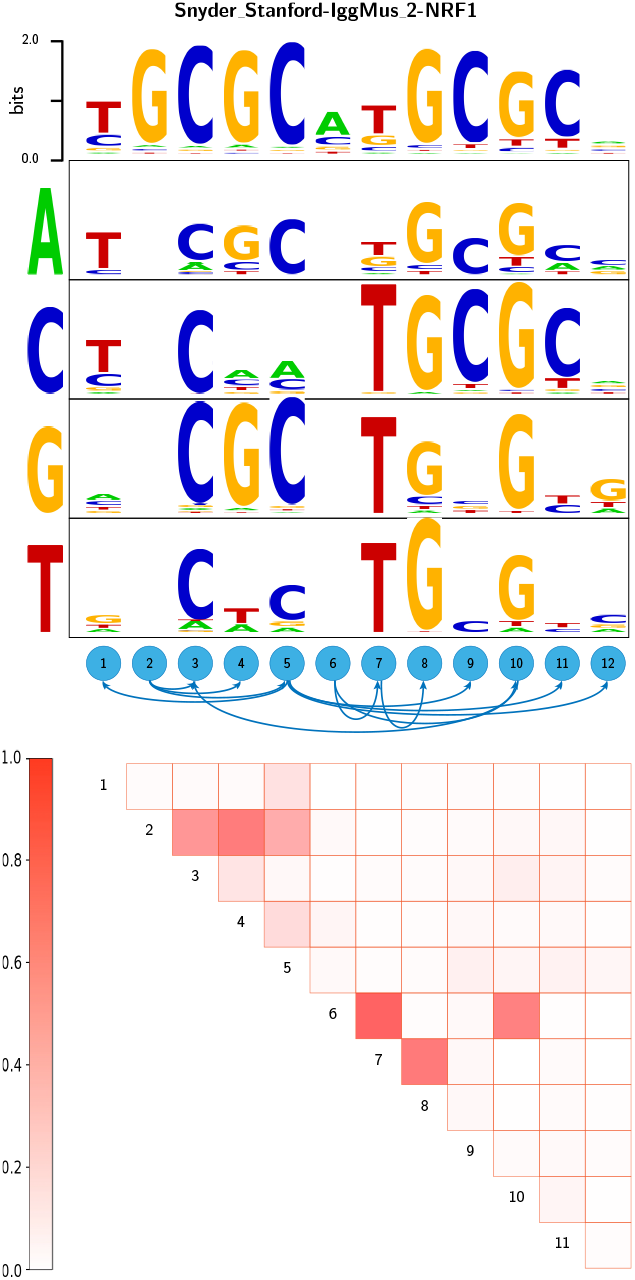
Dilogo for the motif of the TF NRF1. The top row of the dilogo shows the familiar sequence logo representation of the marginal probabilities 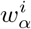 for each of the letters *α* at each position *i*. The posterior probabilities for dependency between each pair of positions are shown in the square lattice at the bottom of the dilogo, with darker red color indicating higher probability of dependence. Above this square lattice a graph with significant pairwise dependencies is shown: an arrow from node *j* to *i* indicates that the probability of a particular letter at *i* depends on the letter appearing at *j*. Finally, for each position *i* that depends on another position *j*, the probabilities *P*(*s*_*i*_*|s*_*j*_) are shown in sequence logo format, with each row corresponding to the identity of the parent letter *s*_*j*_ and each column showing the probabilities *P*(*s*_*i*_*|s*_*j*_) for the child letter *s*_*i*_.

Because it is unwieldy to show the conditional probabilities *P*(*s*_*i*_*|s*_*j*_) for all pairs of positions (*i, j*), we select a set of pairwise dependencies that are jointly consistent with a single factorization of the probability *P*(*S*) as follows. We list all pairwise dependencies *P*(*i, j*), sorted from highest to lowest probability, and go down the list, adding pairwise dependencies as long as the resulting graph does not contain any loops. The resulting graph of dependencies is shown above the square with posterior probabilities. In this example, position 12 depends on position 5, position 11 also depends on position 5, position 10 depends on position 3, and so on.

Finally, for those positions *i* that are dependent on another position *j*, the conditional probabilities *P*(*s*_*i*_*|s*_*j*_) are shown in sequence logo format with one sequence logo (rows in the figure) for each possible state of the parent letter *s*_*j*_ (shown on the left of the figure). For example, in the NRF1 example, the letters at position 3 through 5 depend on the letter at position 2. If position 2 shows a G, positions 3 − 5 are very likely to show the pattern CGC. However, when position 2 shows a T, positions 3 − 5 are most likely to show the pattern CTC.

To enable easy application of DWT models in motif finding we have made a tool-box with software available for motif inference with DWTs, prediction of TFBSs using DWTs, and visualization of DWT models using dilogos. Source code and executables can be downloaded from dwt.unibas.ch.

**Calculating likelihoods of HT-SELEX datasets**

Given a motif model that assigns energies *E*(*s*) to sequence segments *s*, we calculate the likelihood *L*(*E*) of a HT-SELEX dataset as follows. First, for each sequence *S* that occurs in the HT-SELEX data, we calculate a total energy 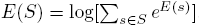. One complication is that the HT-SELEX sequences are all very short, i.e. about 20 nucleotides, such that some motifs can be longer than the input HT-SELEX sequences. To deal with this we padded each HT-SELEX sequence with *l/*2 N nucleotides on both the left and right, where *l* is the motif length, and adapted our sequence scoring to calculate energies for sequence segments containing N nucleotides (see supporting information).

We assume that, in each round of the HT-SELEX experiment, the probability of sampling a sequence *S* is proportional to *e*^*E*(*S*)^. Let *f*_*t*_(*S*) denote the frequency of sequence *S* in the pool of sequences at generation *t* of the HT-SELEX experiment, and let *E*(*S*) denote the total binding energy assigned by the model (either DWT or PSWM) to sequence *S*. Under this model, the probability that a single selected sequence is sequence *S* is given by

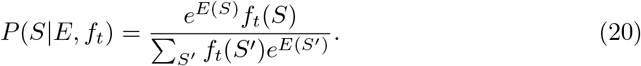

If we denote by *n*_*t*_(*s*) the number of occurrences of sequence *S* at generation *t* in the experiment, then the log-likelihood *L*(*E*) of the entire HT-SELEX data-set, given an energy function *E*, is given by

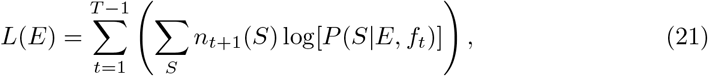

where *T* is the total number of generations in the experiment. This log-likelihood can be compared with the log-likelihood for obtaining the same data by random sampling:

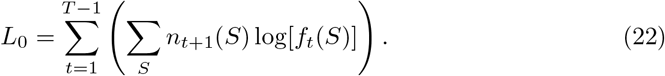

However, when we applied this calculation we find, for almost all corresponding HT-SELEX/ChIP-seq combinations, that the likelihood *L*_0_ is *larger* than the likelihood *L*(*E*), i.e. even for many cases of TFs with well-known motifs. To investigate the origin of this we investigated to what extent the enrichment of sequences from one generation to the next correlates with their predicted energies. In particular, we stratified all sequences into energy bins and calculated the total frequencies *f*_*t*_(*E*) of sequences with predicted energy *E* at each generation *t*. Note that if the probability of a sequence *S* to be selected is proportional to *e*^*E*(*S*)^ then the observed log-enrichment log[*f*_*t*+1_(*E*)*/f*_*t*_(*E*)] should be directly proportional to the energy *E*. However, we observed that, while the log-enrichment generally correlates well with *E*, the slope of the linear relationship is much less than 1, i.e. log[*f*_*t*+1_(*E*)*/f*_*t*_(*E*)] = *βE* + constant, with *β* much smaller than 1. That is, it appears that in HT-SELEX the binding energies vary over a smaller range than predicted by the motif models.

To incorporate this observation, we introduce a ‘temperature’ parameter *β*, assume that the probability of selecting a sequence *S* is proportional to *e*^*βE*(*S*)^, and calculate a log-likelihood *L*(*E, β*) that depends on both the motif model energies *E* and the temperature parameter *β*. It is straightforward to show that the difference *dL*(*E, β*) between the log-likelihood *L*(*E, β*) and the random sampling log-likelihood *L*_0_ can be written as

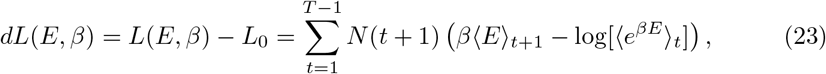

where *N*(*t*) = ∑_*S*_*n*_*S*_ (*t*) is the total number of sequences in generation *t*, ⟨*E*⟩_*t*_ is the average energy of the sequences in generation *t*

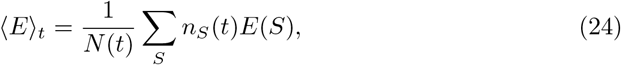

and ⟨*e*^*βE*^⟩_*t*_ is the average selection probability of sequences in generation *t*

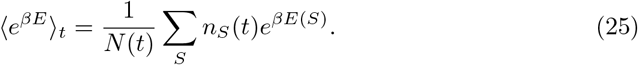

For each PSWM and DWT model, we optimize *β* so as to maximize *dL*(*E, β*) and calculate, as a final performance measure, the log-likelihood difference *dL* per sequence, i.e. *dL/N* with 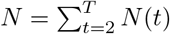.

## Results and Discussion

### DWT models outperform PSWMs and models that only incorporate adjacent dependencies

To compare the performance DWT models with the performance of PSWMs and other motif models, we analyzed a collection of 121 ChIP-seq datasets for 92 different human TFs from the ENCODE consortium [26]. The general setup of our performance comparison is shown in Fig. 2.

We processed each of the ChIP-seq datasets using CRUNCH, an integrated ChIP-seq analysis pipeline that we developed in-house and that includes automated PSWM motif analysis [28]. Full analysis reports on these ChIP-seq datasets as well as links to all the raw ChIP-seq data used are available at crunch.unibas.ch/ENCODE REPORTS/. CRUNCH returns a list of binding peaks, which are typically 100 − 300 base pairs in length, ordered by their significance. For each data-set, we selected the top 1000 binding peaks. The peak sequences were randomly divided into two subsets of 500 sequences, one of which was used as a training set to fit both a PSWM and DWT motif, and one for testing the performance of the fitted motifs. As part of its motif analysis, CRUNCH extracts orthologous sequences from 6 other mammalian species for each peak sequence and multiply aligns these using T-Coffee [29]. The motif finder PhyloGibbs [22] is then run on these alignments to infer PSWM motifs. CRUNCH further refines these motifs on the multiple alignments of the training sequences using MotEvo [30]. For each dataset, we use the top motif returned by CRUNCH as an initial PSWM motif in our analysis and obtained its TFBS predictions on the peak sequences. As an example, Fig. 2b shows the initial PSWM motif inferred for the TF CEBPB.

Using this PSWM as a starting motif we then iteratively fitted a PSWM and a DWT motif on the training sequences (Fig. 2c). The DWT model was fitted using the EM procedure described in the section on motif finding with DWTs above. In order to compare DWTs and PSWMs on equal footing, a PSWM was also fitted on the same training set using the exact same EM procedure.

We then assess the ability of the fitted DWT and PSWM models to explain the ChIP-seq data. In particular, besides the 500 peak sequences of the test set, we created 2000 random decoy sequences that have the same overall dinucleotide frequencies and distribution of lengths as the binding peaks. For each of these 2500 sequences we calculate an overall binding energy *E*(*S*) according to equation (13) using both the PSWM and DWT motifs inferred from the training set. Figure 2d shows the distributions of binding energies that are assigned to the true binding peaks (black) and the decoy sequences (grey) for the fitted PSWM motif, as well as the fitted DWT motif. Comparison of these distributions makes clear that the predicted binding energies of true binding peaks and decoys show a substantially larger separation in the DWT model. Interestingly, this increased separation results mainly from the binding energies of the decoy sequences being more tightly focused at low values. This behavior is observed for a large number of the TFs that we analyzed.

**Fig 2.**
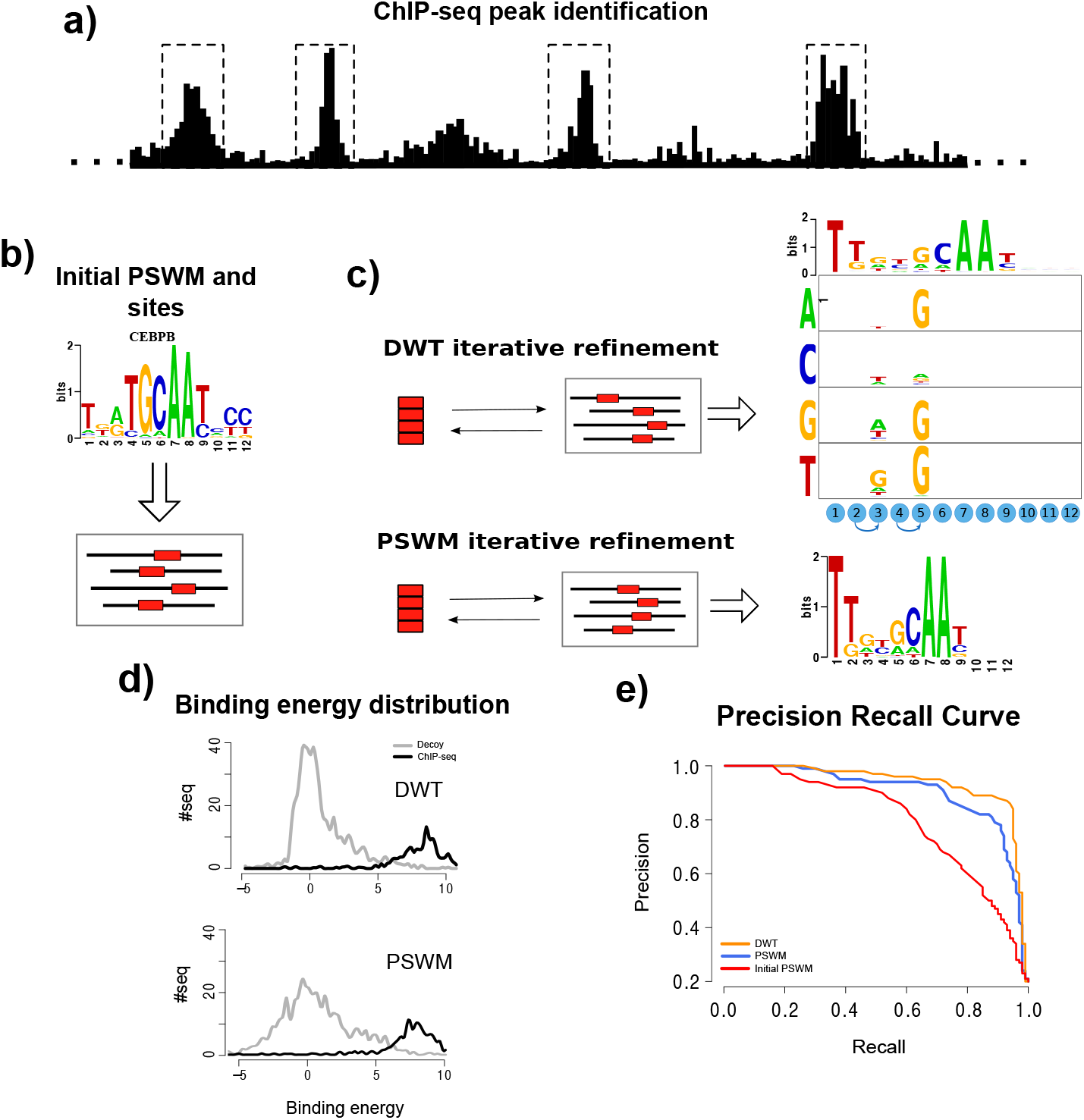
Comparison of DWT and PSWM performance on ChIP-seq data. a) For a given ChIP-seq data-set we use the CRUNCH ChIP-seq analysis pipe-line to identify the top 1000 binding peaks and randomly subdivide these into an training set and a test set of 500 peak sequences each. **b)** Standard PSWM motif finding is used to determine an initial PSWM motif [22, 27]. **c)** Using expectation maximization, a PSWM and a DWT model are fitted on the training data. **d)** Distributions of the predicted binding energies *E*(*S*), under both the DWT and PSWM models, of the 500 peak sequences and a set of 2000 random ‘decoy sequences’ that have the same lengths and dinucleotide composition as the peak sequences. **e)** Precision recall curves demonstrating the ability of the DWT, PSWM, and initial PSWM models to distinguish peak sequences from decoys based on their predicted binding energies.

By systematically varying a cut-off on the binding energy *E*(*S*), we then determine a precision recall curve where, at each cut-off *E*_*c*_, the precision is the fraction of all sequences with *E*(*S*) *> E*_*c*_ that are true binding peaks, and the recall is the fraction of all true binding peaks that have *E*(*S*) *> E*_*c*_. Figure 2e shows the precision recall curves of the original input PSWM, the fitted PSWM, and the DWT model for the TF CEBPB. As a final measure of performance we use the area under the precision-recall curve, which equals the average precision, averaged over all recalls between zero and one.

Figure 3a compares the performance, as measured by average precision, of the DWT and PSWM models on all ENCODE [26] ChIP-seq data-sets that we studied. Remarkably, with the exception of some minor score fluctuations, the DWT model performs at least as well as the PSWM model on all data-sets. This shows that, even though the DWT has no explicit regularization scheme or, in fact, any parameters that need to be set by the user, the model never suffers from over-fitting. Moreover, the DWT model clearly outperforms PSWMs for a substantial fraction of the datasets.

**Fig 3.**
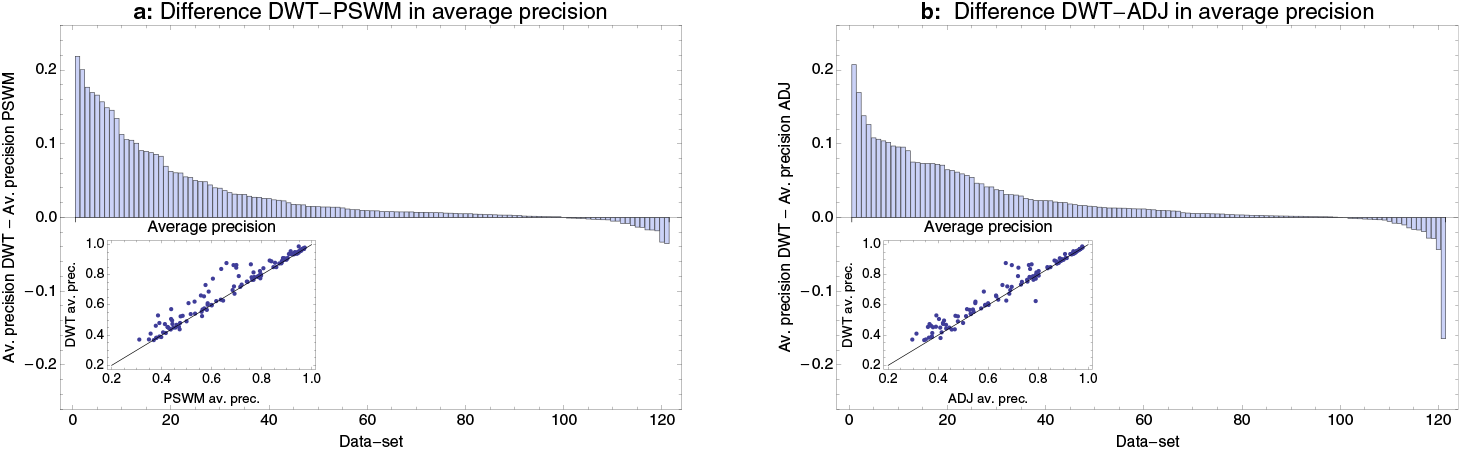
Comparison of the performance of DWT, PSWM, and ADJ Models on the ENCODE ChIP-seq data-sets. **a**: Difference in average precision of the DWT and PSWM models across the 121 ChIP-seq datasets. Datasets are sorted from left to right by the difference in average precision. The inset shows the PSWM average precision (horizontal axis) against the DWT precision (vertical axis), with each dot corresponding to one ChIP-seq dataset, as well as the line *y* = *x*. **b**: As in panel a, but now comparing the average precisions of the DWT model with the ADJ model in which only dependencies between adjacent positions are allowed.

We investigated whether TFs for which the DWT most significantly outperforms the PSWM tend to fall within particular structural families and did not find an any clear association (data not shown). Although it is true that DWTs without any clear pairwise dependencies do not outperform PSWMs, the reverse will not generally hold. That is, the fact that certain positions show clear dependencies does not guarantee that these dependencies will help distinguish binding sites from decoy sequences. Indeed, there are datasets for which DWTs show pairwise dependencies with very high posterior, but where the DWT does not significantly outperform the PSWM. For example, the TF MEF2A shows several pairs of positions with very strong dependency, but the MEF2A DWT does not significantly outperform the corresponding PSWM (see the table with results at http://crunch.unibas.ch/DWT/table.html).

Previous investigations of dependencies between positions in TFBSs have suggested that dependencies between immediately adjacent positions are much more common and significant than dependencies between distal positions [31]. One may thus wonder to what extent the distal dependencies that the DWT infers contribute to the performance of the DWT, or whether a model that uses only dependencies between adjacent positions would perform equally well. As explained in the supporting information, it is straight-forward to adapt the DWT model to only allow dependencies between adjacent positions. We call this version of our motif model the adjacent model (ADJ). We trained and tested the ADJ model in the exact same way as the DWT and PSWM models on all ChIP-seq datasets and Fig. 3b shows the performance comparison between the DWT and ADJ models. We find that the DWT outperforms the ADJ model for more than 80% of the datasets and substantially so for about 25% of the datasets.

Whereas the PSWM never substantially outperformed the DWT (the largest difference in average precision being 3%), there is one dataset for which the ADJ model outperformed the DWT by more than 16% in average precision. This is for ChIP-seq experiment performed in the HeLa cell-line with the chromodomain-like TF CHD2. Notably, the CHD2 TF was also assayed in the GM12878 cell-line, and for this dataset the DWT motif did outperform the ADJ motif. We investigated this case in more detail and found that the DWT had converged to a motif without any significant dependencies, whereas the ADJ had converged to a motif with identical consensus, but with several strong adjacent dependencies. As a test, we reran the DWT motif search on this dataset using the trained ADJ model as a starting motif. We found that the DWT search now converged to a motif that does outperform the ADJ model. That is, there are DWT models that outperform the ADJ for this dataset and the reason the DWT performed poorly was that the motif search happened to have gotten stuck in a poor local optimum.

### DWTs outperform previously proposed motif models that incorporate distal dependencies

Comparing the performance of DWTs with previously proposed approaches is challenging because readily usable software that can be applied to large-scale ChIP-seq results is often not available, and even when software is available it can be challenging to apply it in a manner that allows meaningful performance comparison. Our discussion of the results on the CHD2 dataset underlined that, in order to compare the performance of different motif models, it is essential that all other sources of variability are kept as constant as possible, i.e. not only should we use the exact same train and test data, also the way the motifs are inferred, the way scores of segments are combined to calculate scores of longer sequences, and so on, should be kept as similar as possible. While it was straightforward to accomplish this for comparing our own PSWM, ADJ, and DWT models in the previous section, this is much more challenging when using software from other groups. However, we performed a comparison analysis with two previous methods that allow distal dependencies, for which software was available.

The authors of the FMM method (Sharon *et al.* [18]) and the PIM method (Santolini *et al.* [19]) have not only made software for their motif models available, they also graciously assisted us in adapting their code to allow it to be run in a manner that is as close as possible to the way the DWT model is trained and tested, as detailed in the supporting information.

Figure 4 compares the average precision of the DWT models with those of the PIM and FMM models on the 121 ChIP-seq datasets. We find that, in these tests, the DWT model outperforms the PIM and FMM models on virtually all datasets, and substantially so for a large fraction of the datasets. We want to stress that this does not imply that high-performance PIM and FMM motif models cannot be constructed for these datasets. In our analysis the PIM and FMM motif finders were just run once with default settings and, with appropriate tuning of the parameters, their performance could presumably be substantially improved. However, one of the impediments to the general adoption of more complex motif models has been that running motif inference with these more complex models typically requires expert supervision. One of the main benefits of the DWT model is that it allows robust inference without the need of tuning any parameters.

**Fig 4.**
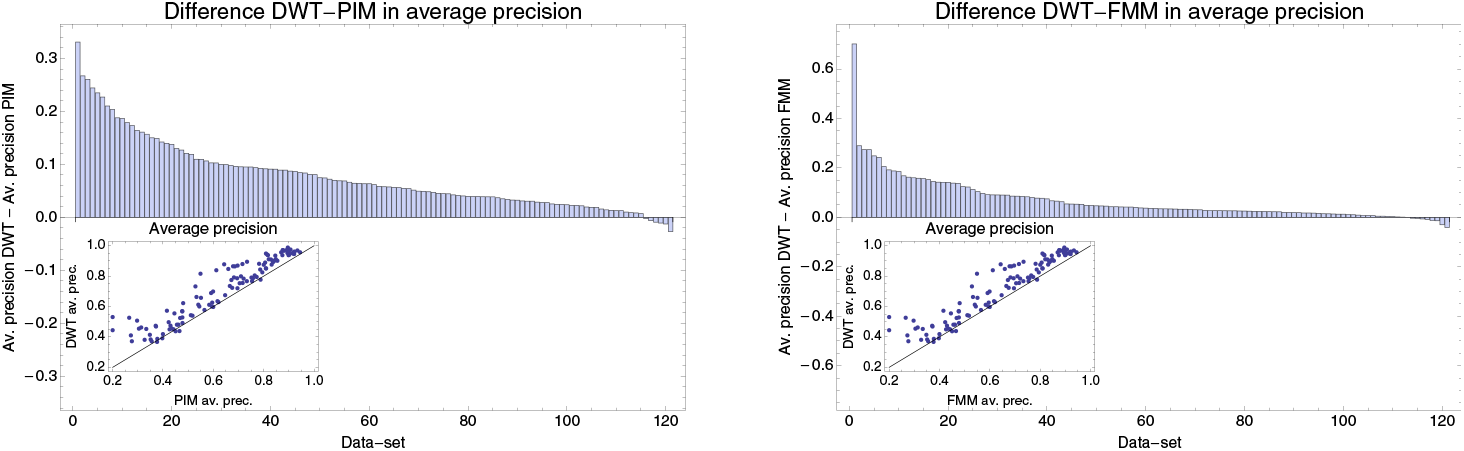
Comparison of the performance of the DWT, PIM [19] and FMM [18] models on the ENCODE ChIP-seq data-sets. **a**: Difference in average precision of the DWT and PIM models across the 121 ChIP-seq datasets. Datasets are sorted from left to right by the difference in average precision. The inset shows the PIM average precision (horizontal axis) against the DWT precision (vertical axis), with each dot corresponding to one ChIP-seq dataset, as well as the line *y* = *x*. **b**: As in panel a, but now comparing the average precisions of the DWT model with the FMM model.

### Pairwise dependencies are enriched at neighboring positions and virtually absent in randomized data

We investigated to what extent pairs of positions that show dependency are restricted to nearest-neighbor interactions. Combining results from all 121 ChIP-seq datasets we calculated the total number of adjacent and and non-adjacent pairs at each posterior probability of dependency. Figure 5 shows the reverse cumulative distributions of the total number of adjacent and distal pairs in our data as a function of their posterior probability. While the absolute number of distal dependencies is consistently above the number of adjacent dependencies at each cut-off, it should be noted that the number of possible distal dependencies is almost 7 times as large as the number of possible adjacent dependencies. Thus, the fraction of adjacent positions that shows dependency is significantly higher than the fraction of distal positions that shows dependency. In summary, adjacent positions are more likely to be dependent than distal positions, although in absolute terms there are more distal dependencies.

**Fig 5.**
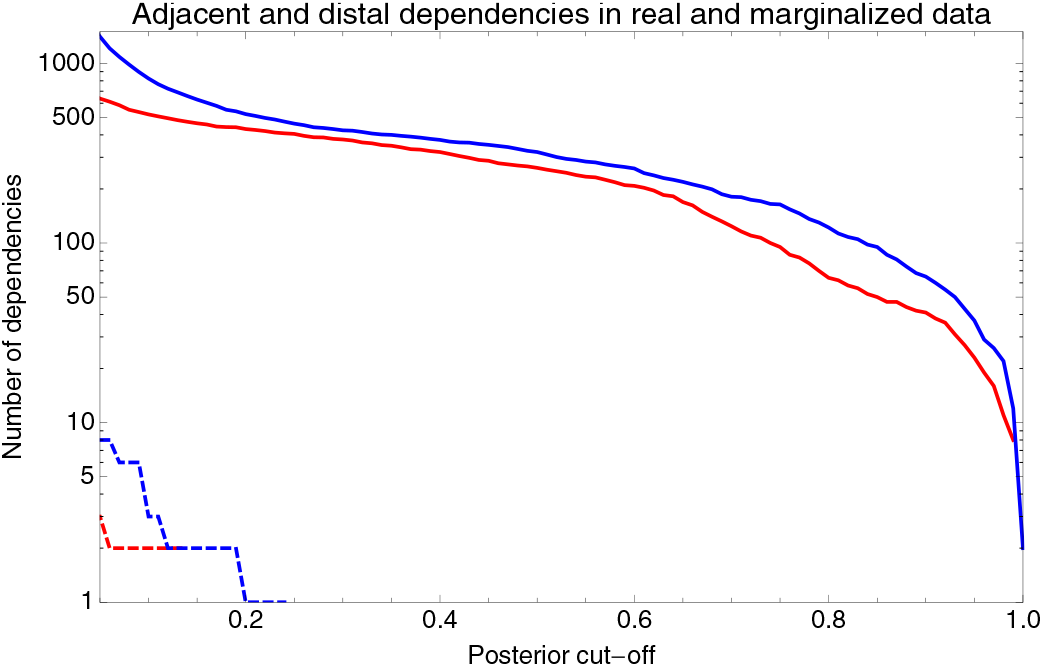
Number of adjacent and distal dependencies as a function of the posterior probability of dependency. The total number of of adjacent (solid red) and distal (solid blue) dependent pairs as a function of a cut-off on the posterior probability of the dependency of the pairs. The dashed lines show the number of adjacent (red) and distal (blue) pairs in randomized data in which DWTs were constructed from sequences sampled from PSWM models.

To confirm the statistical significance of the observed dependencies, we constructed a randomized dataset that should be devoid of dependencies as follows. For each dataset we took the inferred DWT and marginalized it to obtain the corresponding PSWM. When then sampled the same number of binding sites from this PSWM as went into the construction of the DWT, and constructed a new DWT from this set of synthetic binding sites. Finally, we calculated the posterior probabilities of dependency in the set of 121 DWTs so constructed. As shown in Fig. 5, virtually no dependencies appear in this marginalized data, and the dependencies that do appear have low posterior probabilities.

### DWT models trained on ChIP-seq data outperform PSWMs on HT-SELEX data for the same TF

Systematic evolution of ligands by exponential enrichment (SELEX) is a well-established *in vitro* method for studying protein-DNA binding specificity [32]. Starting from a random pool of short DNA (or RNA) segments, the sequences are selected for binding to a DNA protein of interest. The sequences that bound the target are then amplified. This selection and amplification is repeated for multiple rounds to systematically enrich for sequences that strongly bind to the target protein. A high-throughput variant of this method (HT-SELEX), in which the sequences from each round are sequenced using next-generation sequencing was introduced by Jolma *et al.* [33], and has been more recently applied to a large number of human TFs [34]. This HT-SELEX data provides a completely independent dataset for comparing the performance of DWT and PSWM models of TF binding affinities. Moreover, whereas ChIP-seq data arguably probe the *in vivo* binding of a TF in a specific cell type, HT-SELEX probes the binding properties of the DNA binding domain of the TF in an *in vitro* setting. It is thus interesting to investigate whether the DWT outcompetes PSWMs in this *in vitro* setting as well, and to what extent the binding specificities that were inferred from the ChIP-seq data also apply to the HT-SELEX data.

We collected, for each of the TFs assayed in [34], all of our 121 ChIP-seq datasets that were performed with the same (or very similar) TF. In total there were 45 combinations of HT-SELEX/ChIP-seq experiments that were done with the same TF (listed in supplementary table S1). For each combination we then calculated how well the DWT and PSWM models, as inferred from the ChIP-seq dataset, explain the observed HT-SELEX data.

To model the HT-SELEX data we assume that, at each round of the experiment, sequences are selected according to their binding energy to the TF as explained in the materials and methods. As a performance measure of a given motif model, we calculate the average excess of the log-likelihood per selected sequence in each HT-SELEX generation relative to a model which assumes random sampling of sequences. Figure 6 shows the differences in log-likelihood per sequence between the DWT and PSWM models on the 45 combinations of HT-SELEX/ChIP-seq datasets.

For 35 of the 45 combinations, the DWT outperforms the PSWM model on the HT-SELEX data (Fig. 6). Moreover, there is only one example where the PSWM clearly outperforms the DWT model (this example corresponds to the TF IRF4). Note that, although the log-likelihood differences per sequence are typically modest, given the very large number of observations in these HT-SELEX datasets, improvements as small as 0.001 are still highly statistically significant. In summary, for all but one of the TFs, the DWT model that was inferred from ChIP-seq data performs at least as well and often outperforms the PSWM model on HT-SELEX data for the same TF.

**Fig 6.**
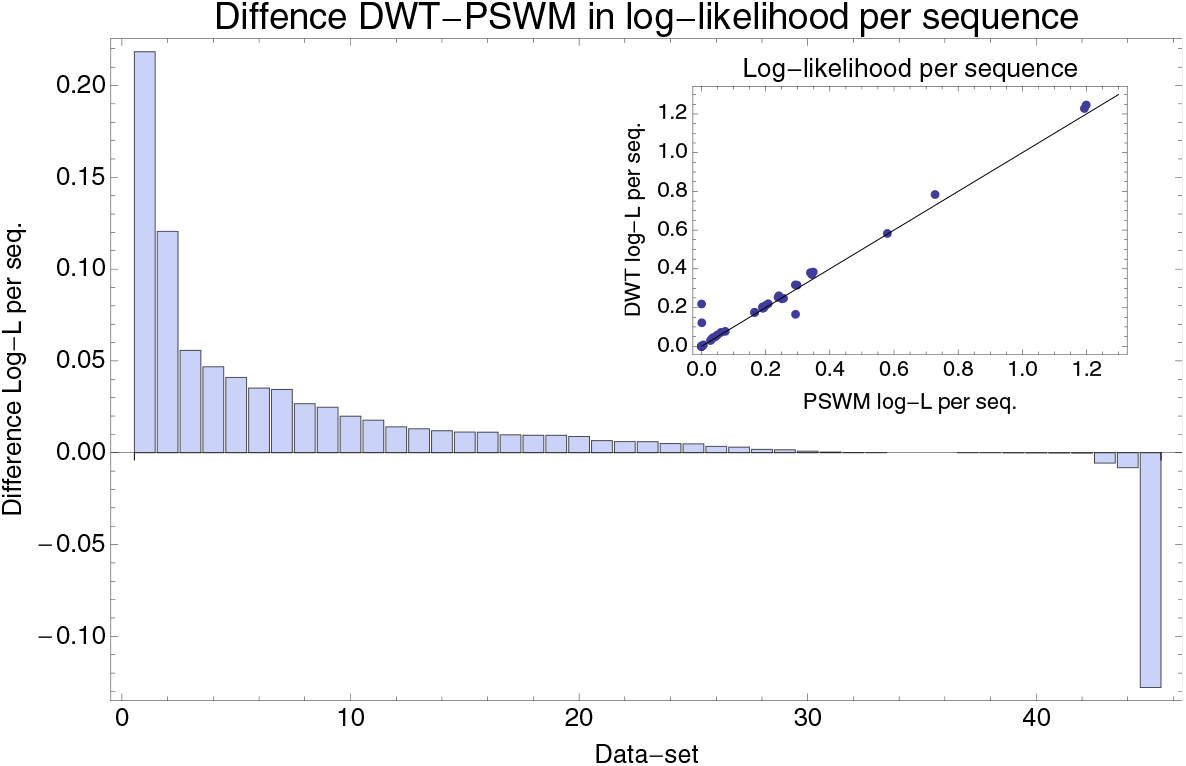
Performance comparison of the DWT and PSWM Models on the HT-SELEX data. Difference in the log-likelihood per sequence between the DWT and PSWM models for each of the 45 corresponding HT-SELEX/ChIP-seq dataset combinations, ordered from left to right by the difference in log-likelihood per sequence. The inset shows the log-likelihood per sequence for the DWT (vertical axis) against the log-likelihood per sequence for the PSWM (horizontal axis), with each dot corresponding to one dataset combination.

## Conclusion

Since its introduction in the early 1980s [35], the PSWM model has become the workhorse for binding site prediction in regulatory genomics. However, as data have accumulated, evidence has been mounting over the last decade that there can be significant dependencies between the nucleotides occurring at different positions of regulatory sites. Consequently, there is a need for extending regulatory motif models to take such dependencies into account. However, in order for such an extension to gain wide acceptance the motif model should be rigorous, flexible, be guaranteed to perform at least as well as PSWMs, and be easy to use. Approaches that have been presented so far have either made unrealistic restrictions on the models, e.g. by demanding that dependencies can only exist between neighboring positions, or they have involved complex *ad hoc* regularization schemes to avoid over-fitting, which make them cumbersome to use in practice.

Here we have presented a new motif model, the dinucleotide weight tensor, that is general in that it allows for dependencies between arbitrary positions in the motif, it is rigorous in that it is derived from first principles within a Bayesian framework, and avoids over-fitting by explicitly marginalizing over all unknown parameters. In particular, because the model has no parameters that the user needs to tune, it can be easily and robustly applied in practice. Indeed, by inferring DWTs on a large set of ChIP-seq datasets, we have shown that DWTs never perform significantly worse than PSWMs and clearly outcompete them in a substantial fraction of the cases. By showing that, for most datasets, DWTs also outperform a model in which only dependencies between adjacent positions are allowed, we further showed that distal dependencies contribute significantly to the performance of the DWTs. We also showed that DWTs outperform two previously proposed methods that incorporate distal dependencies. Notably, while we were finishing this work, a very interesting new approach was proposed by Siebert and Söding [36]. Their motif model is a standard *k*-order Markov model in which each letter depends on the (*k* − 1) previous letters in the site, but a new way for controlling over-fitting is proposed, in which the marginals at lower orders are used a priors for the conditional probabilities at higher orders, and very robust performance of this method is proposed. Interestingly, it would be straightforward to combine this method of setting priors for conditional probabilities with the DWT’s method for summing over possible spanning trees, and this would be an interesting direction to explore for future work.

The fact that DWT models inferred from ChIP-seq data also outperform PSWMs on HT-SELEX data, suggests that the dependencies captured by the DWT reflect something in the biophysics of the interaction between the DNA binding domain of the TF and the DNA sequence of the site. Our observation that, while significant dependencies occur between distal positions, interactions between neighboring positions are the most common, is also consistent with this interpretation. Another interesting area for future research is to investigate the possible structural and biophysical basis for the observed direct dependencies. However, we should note that, in spite of investing considerable efforts ourselves in analyzing whether the occurrence of dependencies can be related to structural features of the TFs, or to the way that they interact with the DNA, we have so far not been able to uncover any consistent biophysical interpretation of the observed dependencies. It is conceivable that there is no simple biophysical interpretation to the direct dependencies. For example, inspection of some of the DWT models suggests that dependencies often cause combinations of deleterious mutations to reduce the binding energy less than predicted by the PSWM model and this might be a global effect that is spread across many dependencies, rather than reflecting particular structural features of the TF-DNA interaction.

Our analysis has also shown that, notwithstanding the fact that DWTs strongly outperform PSWMs for some TFs, for the majority of TFs the improvement that the DWT provides is rather modest. This highlights that, for many TFs, PSWMs are sufficiently accurate for TFBS prediction, and few significant dependencies exist. Consequently, robust practical application of more complex motif models requires strong safe-guards against over-fitting, i.e. because for many TFs there will simply not be many strong dependencies. This is arguably the biggest advantage of the DWT models presented here: DWTs have no parameters to tune, do not overfit, and automatically reduce to a PSWM model when no significant dependencies exist. We believe that these properties make DWTs especially attractive for adopting in practical settings and we hope that many researchers can be convinced to start using DWT models in their motif finding and TFBS prediction.

## Acknowledgments

SO thanks Lukas Burger for help with the Bayesian model and its implementation, and Peter Pemberton-Ross and Stephanie Bishop for help with the writing of the manuscript. This work was supported by SystemsX.ch through the CellPlasticity project grant.

## Supporting text S1

Supporting mathematical derivations and a table of all HT-SELEX/ChIP-seq combinations that we analyzed

